# EphrinB2 knockdown in cervical spinal cord preserves diaphragm innervation in a mutant SOD1 mouse model of ALS

**DOI:** 10.1101/2023.05.10.538887

**Authors:** Mark W. Urban, Brittany A. Charsar, Nicolette M. Heinsinger, Shashirekha S. Markandaiah, Lindsay Sprimont, Wei Zhou, Eric V. Brown, Nathan T. Henderson, Samantha J. Thomas, Biswarup Ghosh, Rachel E. Cain, Davide Trotti, Piera Pasinelli, Megan C. Wright, Matthew B. Dalva, Angelo C. Lepore

## Abstract

Amyotrophic lateral sclerosis (ALS) is a neurodegenerative disease characterized by motor neuron loss. Importantly, non-neuronal cell types such as astrocytes also play significant roles in disease pathogenesis. However, mechanisms of astrocyte contribution to ALS remain incompletely understood. Astrocyte involvement suggests that transcellular signaling may play a role in disease. We examined contribution of transmembrane signaling molecule ephrinB2 to ALS pathogenesis, in particular its role in driving motor neuron damage by spinal cord astrocytes. In symptomatic SOD1^G93A^ mice (a well-established ALS model), ephrinB2 expression was dramatically increased in ventral horn astrocytes. Reducing ephrinB2 in the cervical spinal cord ventral horn via viral-mediated shRNA delivery reduced motor neuron loss and preserved respiratory function by maintaining phrenic motor neuron innervation of diaphragm. EphrinB2 expression was also elevated in human ALS spinal cord. These findings implicate ephrinB2 upregulation as both a transcellular signaling mechanism in mutant SOD1-associated ALS and a promising therapeutic target.

## INTRODUCTION

Astrocytes are glial cells that play critical roles in central nervous system (CNS) function and dysfunction, including in neurodegenerative diseases such as ALS ^1^. In ALS, loss of upper motor neurons (MNs) in the brain and lower MNs of spinal cord and brainstem results in progressive muscle paralysis and ultimately in death, usually in only 2-5 years after diagnosis ^2^. The majority of ALS cases are sporadic, while 10% are of the familial form; these familial cases are linked to a variety of genes such as Cu/Zn superoxide dismutase 1 (SOD1) ^3^, TAR DNA-binding protein 43 (TDP-43) ^4^, C9orf72 hexanucleotide repeat expansion ^5,6^ and others.

While ALS is characterized primarily by MN degeneration, studies with human ALS tissue and experiments in animal and in vitro models of ALS demonstrate that cellular abnormalities are not limited to MNs ^7^. In particular, non-neuronal cell types such as astrocytes play significant roles in disease pathogenesis. Findings suggest that transcellular signaling between astrocytes and MNs may represent an important regulatory node for MN survival and disease progression in ALS ^8^. However, the mechanistic contributions of astrocytes to ALS remain incompletely understood, hampering development of effective therapies for targeting this cell population and for treating the disease.

One family of proteins linked to both transcellular signaling and ALS are the erythropoietin-producing human hepatocellular receptors (Ephs) and the Eph receptor-interacting proteins (ephrins) ^9^. Ephs are transmembrane signaling molecules and the largest known family of receptor tyrosine kinases in the mammalian genome. Ephs bind to and are activated by ephrins, which are either glycosylphosphatidylinositol-linked (ephrin-As) or are transmembrane proteins (ephrin-Bs) also capable of signaling ^10,11^. Eph-ephrin transcellular signaling regulates many events in the developing and mature nervous system that are mediated by cell contact dependent mechanisms, including: dendritic spine formation, dendritic filopodia dependent synaptogenesis, axon guidance, control of synapse maintenance and density, and synaptic localization of glutamate receptor subunits ^10,11^. Eph-ephrin signaling is an important mediator of signaling between neurons and non-neuronal cells in the nervous system. Neuronal EphA binding to glial ephrin plays an important role in the morphogenesis of dendritic spines ^12^. In the peripheral nervous system, axons expressing EphA are guided to their correct target via ephrin-Bs expressed in the limb bud. Moreover, during development EphA4 is expressed by MNs undergoing programed cell death ^13^, while blockade of EphA4 signaling can limit cell death in models of stroke ^14,15^. Thus, Eph-ephrin transcellular signaling is a potent modulator of neuronal function and survival.

In the mature CNS, dysregulation of Eph and ephrin signaling has been linked to a number of neurodegenerative diseases, including ALS. Expression of EphA4 in MNs significantly contributes to MN degeneration and overall disease pathogenesis in both rodent and zebrafish animal models of ALS ^16^, while reduction of ephrinA5 worsens disease outcome in an ALS mouse model ^17^.

Furthermore, increased EphA4 expression levels and EphA4 signaling capacity correlate with the degree of human ALS disease severity ^16^. While antisense oligonucleotide ^18^ or ubiquitous genetic knockdown ^19^ of EphA4 in ALS mouse models does not affect disease phenotype, inhibition of EphA4 signaling using EphA4-Fc partially preserves motor function and MN-specific genetic knockdown delays symptomatic onset and protect MNs in ALS mice ^20^.

In search of new targets to modulate Eph-ephrin signaling, we chose to explore ephrinB2 given previous work showing its role in astrocytes in the disease pathology of other neurological conditions such as traumatic spinal cord injury ^21,22^. We find that ephrinB2 expression in ventral horn astrocytes increases with disease progression in the SOD1^G93A^ mouse model of ALS. Patients ultimately succumb to ALS because of respiratory compromise due in part to loss of respiratory phrenic MNs (PhMNs) that innervate the diaphragm ^23^. We therefore tested in the current study viral vector-based small hairpin RNAs (shRNA) knockdown of ephrinB2 ^24^ in the ventral horn of SOD1^G93A^ cervical spinal cord ^25^. We evaluated *in vivo* effects on key outcomes associated with human ALS, including protection of cervical MNs ^26–28^, maintenance of diaphragm function and innervation by PhMNs ^25,29,30^, and overall phenotypic disease extension ^31^. Collectively, data from our study provides insights into both disease mechanisms governing MN loss in mutant SOD1 ALS and a potential therapeutic target.

## RESULTS

### Increase in ventral horn ephrinB2 with disease progression

Eph receptor signaling has been implicated in ALS ^16,32^; however, the involvement of specific ephrin ligands in disease remains unresolved. To begin to address this question, we assessed ephrinB2 expression over the course of disease in SOD1^G93A^ mice at pre-symptomatic (60 days), symptomatic (120 days) and endstage time points using ephrinB2 immunohistochemistry (IHC). We focused in particular on the cervical ventral horn, as it is the location of PhMNs critical to maintaining diaphragm function ^33^. In age-matched wild-type (WT) littermates, ephrinB2 was expressed at relatively low levels. Compared to WT controls (Fig 1a), there was pronounced up-regulation of ephrinB2 in ventral horn even at the late pre-symptomatic (60 day) time point (Fig 1b). EphrinB2 expression dramatically increased over disease course in SOD1^G93A^ mice as seen at the symptomatic (120 day) time point (Fig1 c) and at disease endstage (Fig1 d). Quantification of ephrinB2 expression in the cervical spinal cord ventral horn showed an increase in expression at 60 days (18.58 ± 8.42 a.u. fold increase; WT vs. 60d: p = 0.048), 120 days (41.83 ± 26.67 a.u. fold increase; WT vs. 120d: p = 0.21; 60d vs. 120d: p = 0.32) and endstage (63.42 ± 10.99 a.u. fold increase; WT vs. endstage: p = 0.015; 60d vs. endstage: p = 0.017; 120d vs. endstage: p = 0.3) compared to WT age-matched controls (1.00 ± 0.40 a.u.) (Fig 1e; n = 3-4 mice per group). Compared to WT control, ephrinB2 expression was also significantly increased at endstage in the thoracic (Fig 1f, g) and lumbar (Fig 1h-k) ventral horn. Increases in ephrinB2 expression were localized to spinal cord gray matter. Higher magnification imaging from lumbar spinal cord revealed that the vast majority of ephrinB2-expressing cells within the ventral horn displayed an astrocyte-like morphology (Fig 1j-k). These data indicate that ephrinB2 expression is upregulated in SOD1^G93A^ mice and suggest that increases in ephrinB2 expression might be localized to glia.

**Figure 1:**
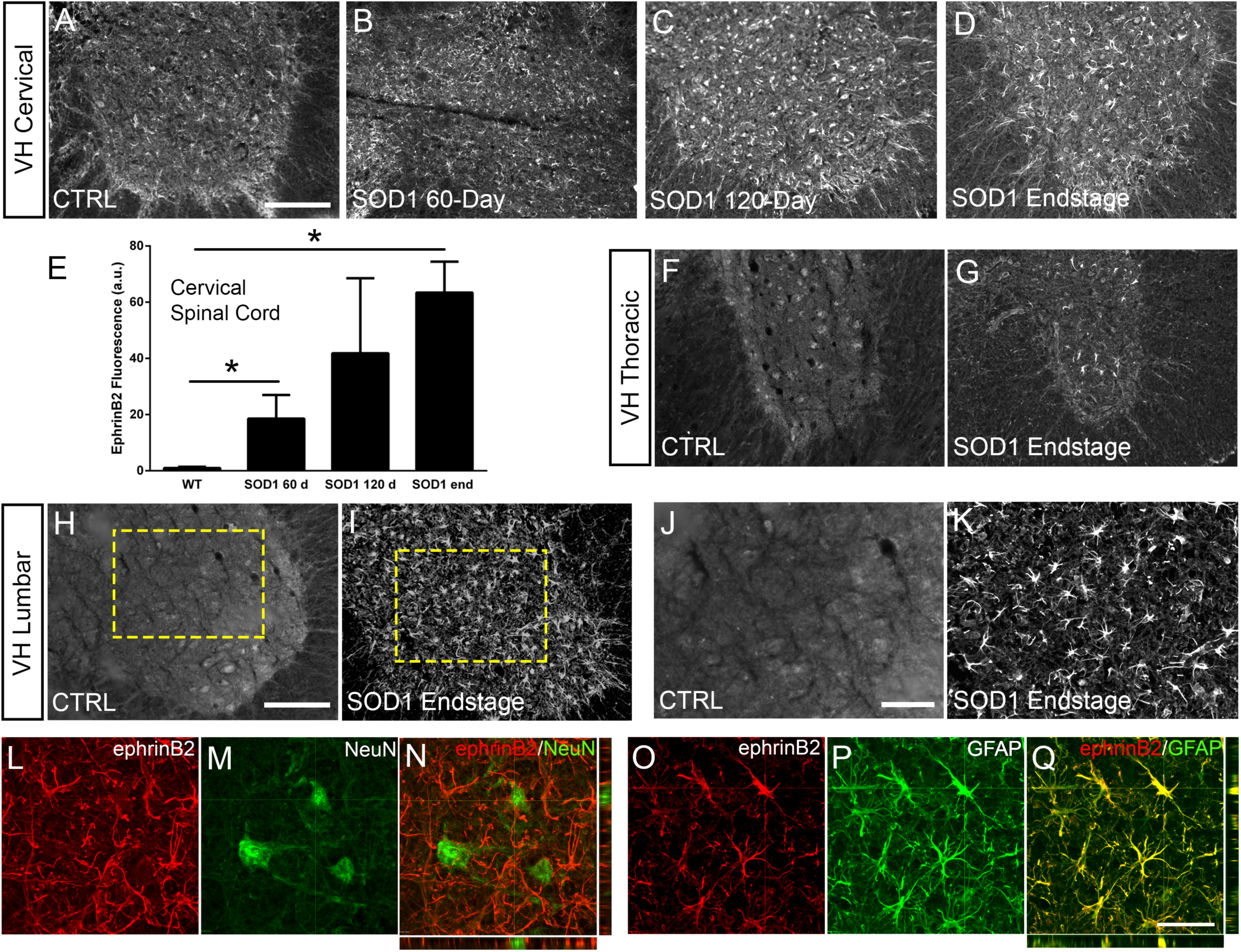
EphrinB2 expression was increased in ventral horn astrocytes. SOD1^G93A^ mouse ventral horn cervical spinal cord tissue immuinostained for ephrinB2 at 60 days (**b**), 120 days (**c**), endstage (**d**), and WT mouse age-matched control (**a**), scale bar: 200 µm. Quantification of ephrinB2 expression within the ventral horn shows a progressive increase in expression over time compared to WT controls (**e**). Endstage ephrinB2 expression in the thoracic (**f**, **g**) and lumbar (**h**-**k**) regions; scale bar: 200 µm, 100 µm, respectively. Endstage SOD1^G93A^ mouse cervical spinal cord tissue co- immuostained for ephrinB2 (**l**, **n**, **o**, **q**) and neuronal and astrocyte lineage-specific markers NeuN (**m-n**) and GFAP (**p-q**), respectively; scale bar: 30 µm. Analysis in panels A-K: n = 3-4 mice per genotype and per time point; 1-2 females and 2 males per condition. Analysis in panels L-O: n = 3 mice per genotype and per time point; 1 female and 2 males per condition.

### EphrinB2 was upregulated in ventral horn astrocytes

We next asked whether ephrinB2 was expressed in astrocytes. Expression of ephrinB2 was determined in neurons and astrocytes within ventral horn at disease endstage using double-IHC for ephrinB2 along with lineage-specific antibodies for reactive astrocytes (GFAP: glial fibrillary acidic protein) and for neurons (NeuN: neuronal nuclear protein) ^34^. EphrinB2 upregulation was localized to GFAP-expressing astrocytes (Fig 1o-q) and was not co-localized to NeuN-expressing neurons (Fig 1l- n) (n = 3 mice). Thus, ephrinB2 expression is dramatically and selectively increased in reactive astrocytes of the SOD1^G93A^ mouse spinal cord in areas of MN loss.

### EphrinB2 knockdown in astrocytes of cervical ventral horn

Given that both ALS patients ^23^ and mutant SOD1 rodents ^35^ succumb to disease due in part to diaphragmatic respiratory compromise, we next sought to focally reduce ephrinB2 expression in astrocytes in the region of the spinal cord containing respiratory PhMNs. To begin to test whether the increased expression of ephrinB2 might impact disease progression, we injected 60 day old SOD1^G93A^ mice with either lentivirus-GFP control vector or lentivirus that transduces an ephrinB2 shRNA expression cassette ^24^. Virus was injected bilaterally into the ventral horn at 6 sites throughout the C3-C5 region to bilaterally target the region of the spinal cord containing the diaphragmatic respiratory PhMN pool (Fig 2a-b) ^25^. We have shown previously that this shRNA construct selectively targets ephrinB2 and that knockdown effects of the shRNA on ephrinB2 levels are rescued by expression of a shRNA-insensitive version of ephrinB2 ^24^.

**Figure 2:**
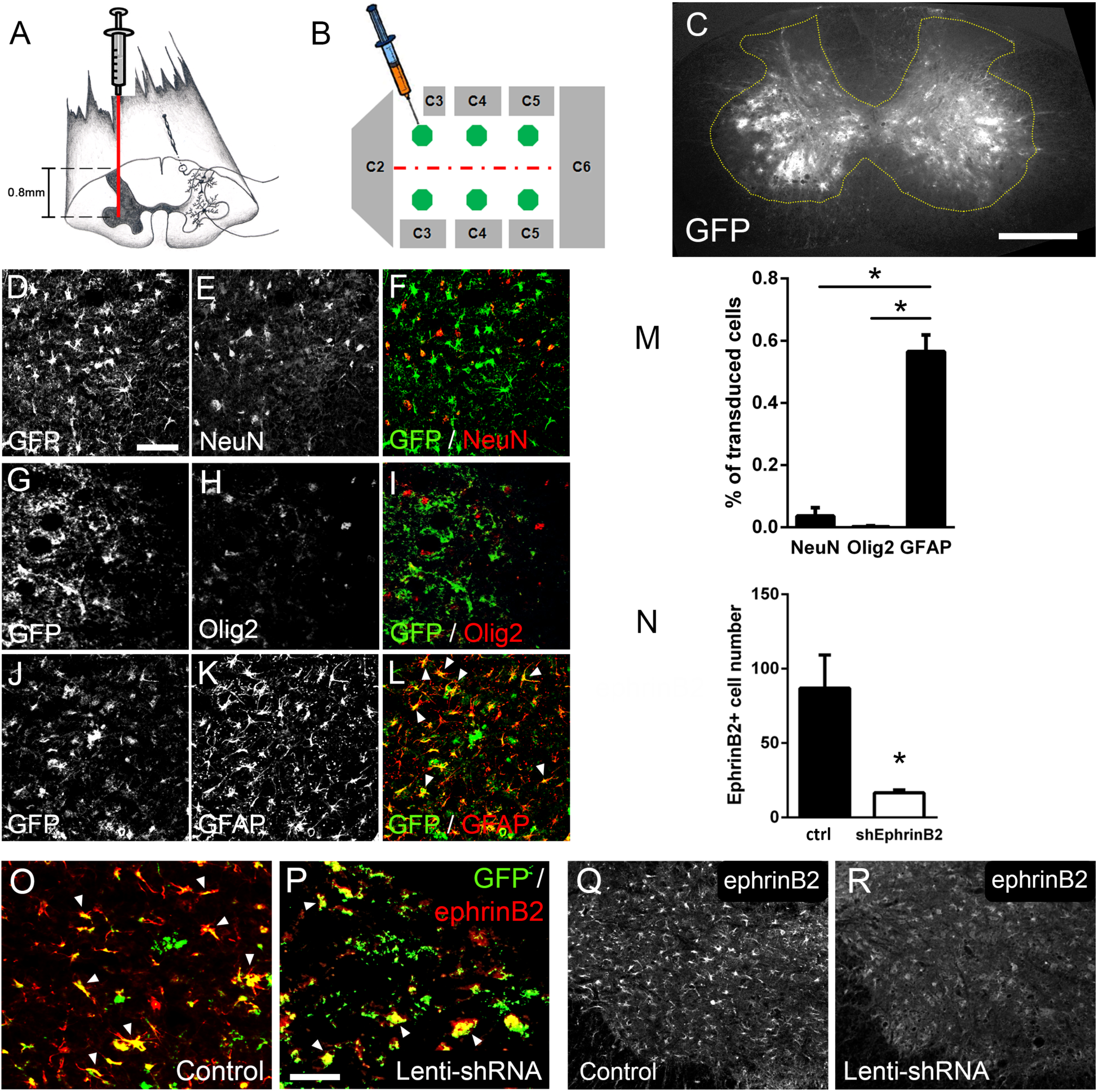
Lenti-shRNA injection reduced ephrinB2 expression in cervical spinal cord astrocytes. We injected 60 day old SOD1^G93A^ mice with Lenti-GFP control or Lenti-shRNA-ephrinB2 into cervical ventral horn (**a**). Injections were made bilaterally into 6 sites throughout C3-C5 to target the PhMN pool (**b**). 30 µm transverse tissue sections of the cervical spinal cord show robust expression of the GFP reporter bilaterally within the ventral horn (**c**); scale bar: 500 µm. Cell lineage of viral transduction was assessed using three markers: neuronal marker NeuN, oligodendrocyte lineage marker Olig2, and astrocyte marker GFAP. Spinal cord tissue collected from SOD1^G93A^ injected with Lenti-GFP vector was sectioned at 30 µm and immunostained for NeuN (**d**-**f**), Olig2 (**g**-**i**) and GFAP (**j**-**l**); scale bar: 150 µm. Quantification of transduction lineage was assessed by counting the total numbers of GFP+ cells that were co-labeled with each lineage-specific marker and expressing this as a percentage of the total number of GFP+ cells (**m**). We assessed the amount of knockdown achieved by the Lenti-shRNA-ephrinB2 vector by immunostaining endstage SOD1^G93A^ cervical spinal cord tissue with an anti-ephrinB2 antibody in both Lenti-GFP control (**o, q**) and Lenti-shRNA-ephrinB2 (**p, r**) tissue; scale bar: 100 µm. Knockdown was quantified by counting the total number of GFP+ cells expressing ephrinB2+ within the cervical ventral horn (**n**). Analyses in all panels: n = 3 mice per condition; 1 female and 2 males per condition.

We first determined whether our knockdown approach efficiently transduced astrocytes and reduced ephrinB2 expression in spinal cord astrocytes. Transverse sections of endstage SOD1^G93A^ mouse cervical spinal cord show robust expression of the GFP reporter bilaterally within ventral horn following intrapsinal injection (Fig 2c). To evaluate cell lineage of viral transduction, we performed IHC on lenti-GFP transduced spinal cord tissue at disease endstage. The majority of GFP-expressing cells in ventral horn were GFAP+ reactive astrocytes; these GFAP+/GFP+ astrocytes also expressed high levels of ephrinB2, demonstrating that the lentiviral constructs targeted reactive astrocytes that included those with upregulated ephrinB2 expression (Fig 2j-l). On the contrary, there was little-to-no co-labeling of the GFP reporter with NeuN+ neurons (Fig 2d-f) or cells positive for oligodendrocyte transcription factor 2 (Olig2) (Fig 2g-i), demonstrating that the injected viral constructs did not target a large portion of neurons or cells of the oligodendrocyte lineage within the ventral horn. We quantified the percentage of transduced GFP+ cells that co-labeled with GFAP, NeuN or Olig2 and found that the majority of transduced cells were astrocytes (NeuN: 3.54 ± 1.09 % labeled cells, n = 3; Olig2: 0.18 ± 0.18 % labeled cells, n = 3; GFAP: 56.53 ± 5.30 % labeled cells, n = 3 mice; GFAP vs. NeuN: p = 0.023; GFAP vs. Olig2: p = 0.018) (Fig 2m). We next determined whether the lenti-shRNA vector effectively reduced ephrinB2 expression in ventral horn astrocytes. Compared to lenti-GFP control (Fig 2o, q), the lenti-shRNA (Fig 2p, r) reduced ephrinB2 expression by approximately a factor of 5 (Lenti-GFP: 86.92 ± 22.35 GFP+/ephrinB2+ cells, n = 3 mice; Lenti-shRNA: 16.67 ± 1.76 GFP+/ephrinB2+ cells, n = 3 mice mice; t-test, p = 0.035) (Fig 2n). Together, these results show that viral transduction was anatomically-targeted to the cervical ventral horn, was relatively-specific to the astrocyte lineage, and was able to significantly reduce ephrinB2 expression levels within the C3-C5 ventral horn of SOD1^G93A^ mice.

### Protection of MNs in the cervical spinal cord

Loss of motor neurons (MNs) in spinal cord is a hallmark of ALS. To determine whether knockdown of ephrinB2 in astrocytes might impact MN survival selectively in the region of ephrinB2 knockdown, we quantified MN somata within the C3-5 spinal cord. Using cresyl violet staining of transverse cervical spinal cord sections, the number of neurons with a somal diameter greater than 20 µm and with an identifiable nucleolus was determined (MNs, Fig 3a) ^25^. In C3, C4 and C5 following transduction of Lenti-shRNA-ephrinB2 (Fig 3d), there was a significantly greater number of MNs within the ventral horn compared to Lenti-GFP controls (Fig 3c) (Lenti-GFP: 266.4 ± 19.46 MNs/µm^2^, n = 4 mice; Lenti- shRNA-ephrinB2: 344.3 ± 6.31 MNs/µm^2^, n = 4 mice; p = 0.019, t-test) (Fig 3b). These data suggest that knockdown of ephrin-B2 can increase survival of MNs in a mutant SOD1 model of ALS.

**Figure 3:**
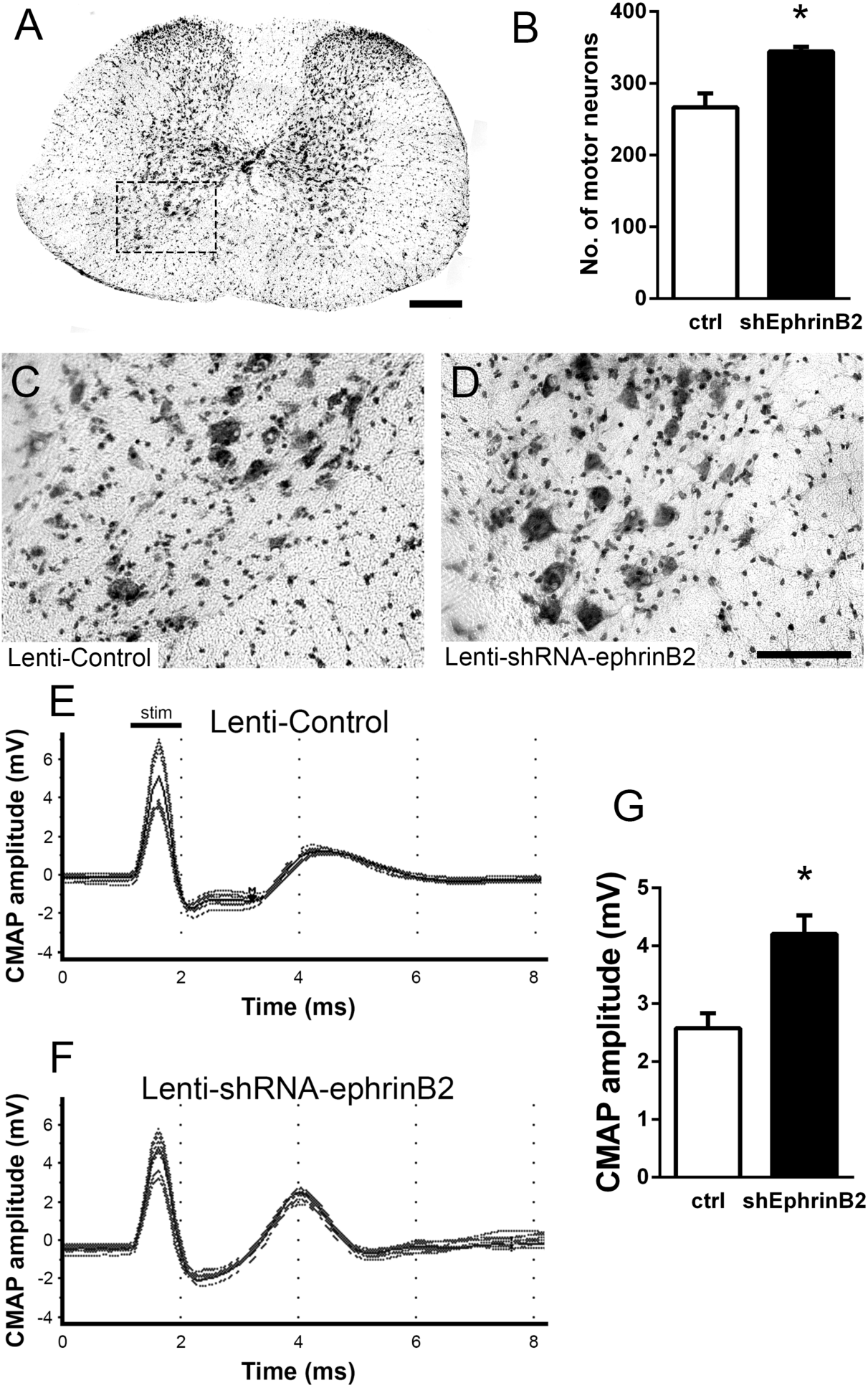
EphrinB2 knockdown protected cervical spinal cord motor neurons and preserved functional innervation of the diaphragm in SOD1^G93A^ mice. SOD1^G93A^ mice injected with Lenti- GFP control or Lenti-shRNA-ephrinB2 were cresyl violet stained and MN counts were performed at 117 days of age. 30 µm transverse cervical spinal cord tissue sections were stained with cresyl violet (**a**); scale bar: 250 µm. The dotted box outlines the ventral horn and area of the image shown in (**c**). MN populations within the ventral horn were quantified (**b**). Representative images show a greater loss of MNs in Lenti-Control (**c**) compared to the Lenti-shRNA-ephrinB2 (**d**) group; scale bar: 100 µm. SOD1^G93A^ mice injected with Lenti-GFP control or Lenti-shRNA-ephrinB2 were assessed in vivo for PhMN-diaphragm innervation by electrophysiological analysis at 117 days of age. CMAP amplitudes were recorded from each hemi-diaphragm following ipsilateral phrenic nerve stimulation. Representative traces of Lenti-GFP (**e**) and Lenti-shRNA-ephrinB2 (**f**) recordings show a larger in CMAP amplitude in the Lenti-shRNA-ephrinB2 group. Quantification of maximal CMAP amplitude shows significant preservation in the Lenti-shRNA-ephrinB2 treated group compared to control (**g**). Analysis in panels A-D: n = 4 mice per condition; 2 females and 2 males per condition. Analysis in panels E-G: n = 4 mice per genotype and per time point; 2 females and 2 males per condition.

### Preservation of diaphragm function

Patients ultimately succumb to ALS because of respiratory compromise due significantly in part to loss of PhMNs that innervate diaphragm, the primary muscle of inspiration ^23^. To evaluate whether ephrinB2 knockdown in astrocytes focally within the PhMN pool impacts respiratory neural circuitry, we determined effects on both PhMN innervation of diaphragm using morphological assessment ^26–28^ and preservation of diaphragm function using *in vivo* electrophysiological measurements ^25,26,29,30^. In anesthetized mice, we recorded compound muscle action potential (CMAP) amplitudes from each hemi-diaphragm following supramaximal stimulation of the ipsilateral phrenic nerve, an electrophysiological assay of functional diaphragm innervation by PhMNs. We performed these experiments in SOD1^G93A^ mice at 117 days of age, a time point following the beginnings of forelimb motor dysfunction in the vast majority of animals but prior to endstage. Quantification of CMAP amplitude showed a 61% larger amplitude for the lenti-shRNA group (Fig 3f) compared to lenti-GFP (Fig 3e, g), demonstrating that ephrinB2 knockdown in cervical ventral horn resulted in significant preservation of functional diaphragm innervation (Lenti-GFP: 2.58 ± 0.26 mV, n = 4 mice; Lenti-shRNA: 4.20 ± 0.32 mV, n = 4 mice; t-test, p = 0.0075). These data indicate that region-specific knockdown of ephrinB2 was able to generate functional rescue appropriate for the location targeted.

### Effects on disease onset, disease duration or animal survival

We chose to perform anatomically-targeted shRNA delivery to only the ventral horn of the cervical (C3-C5) spinal cord in order to specifically target the critically-important phrenic nucleus and to use this motor circuit as a model system to examine the impact of knocking down astrocyte ephrinB2 expression on PhMN degeneration and diaphragm innervation. As expected, given that injections were delivered only to levels C3-5, ephrinB2 knockdown in astrocytes had no impact on overall disease phenotype, including limb motor function, disease onset and progression, and animal survival, as assessed by a battery of established measurements ^25,29–31^. EphrinB2 knockdown did not affect weight loss at any age tested (F (1, 18) = 0.17, p = 0.69) (Fig 4a; n = 8-10 mice per group). Additionally, overall disease onset as determined by the timing of weight loss onset was unaffected, with both the lenti-GFP and lenti-shRNA groups showing similar onset as determined by Kaplan-Meier analysis (Lenti-GFP: 123.5 days; Lenti-shRNA-ephrinB2 125.0 days, chi square: 0.017, p = 0.90, Gehan-Breslow-Wilcoxon test; n = 8-9 mice per group) (Fig 4b). Furthermore, there were no differences between the two groups in either hindlimb (F (1, 18) = 0.48, p = 0.50, ANOVA; n = 9-10 mice per group) (Fig 4c) or forelimb (F (1, 18) = 0.95, p = 0.34, ANOVA; n = 9-10 mice per group) (Fig 4e) grip strength decline. We also used these grip strength measurements to calculate hindlimb and forelimb disease onsets. We calculated onset individually for each animal as the age with a 10% decline in grip strength compared to the maximum strength for those limbs in the same animal ^29,31^. EphrinB2 knockdown had no effect on either hindlimb onset (Lenti-GFP: 90.5 days; Lenti-shRNA- ephrinB2 108.0 days, chi square: 2.92, p = 0.09, Gehan-Breslow-Wilcoxon test; n = 9-10 mice per group) (Fig 4d) or forelimb onset (Lenti-GFP: 116.5 days; Lenti-shRNA-ephrinB2 111.0 days, chi square: 0.13, p = 0.72, Gehan-Breslow-Wilcoxon test; n = 9-10 mice per group) (Fig 4f). Given that previous work showed that astrocytes contribute to disease progression in mutant SOD1 rodents post-disease onset ^36^, we examined whether ephrinB2 knockdown in astrocytes extended disease duration. Compared to lenti-GFP control, lenti-shRNA had no effect on disease duration as measured by the time from: weight onset to endstage (Lenti-GFP: 7.90 ± 1.110 days, n = 10 mice; Lenti-shRNA-ephrinB2: 11.38 ± 1.963 days, p = 0.13, unpaired t-test; n = 8 mice) (Fig 4g); hindlimb disease onset to endstage (Lenti-GFP: 41.90 ± 6.63 days, n = 10 mice; Lenti-shRNA-ephrinB2: 29.44 ± 6.34 days, n= 9 mice, p = 0.19, unpaired t-test) (Fig 4h); forelimb disease onset to endstage (Lenti-GFP: 25.10 ± 6.48 days, n = 10 mice; Lenti-shRNA-ephrinB2: 27.11 ± 8.92 days, n = 9 mice, p = 0.86, unpaired t-test) (Fig 4i); or hindlimb disease onset to forelimb disease onset (Lenti-GFP: 16.80 ± 4.14 days, n = 10 mice; Lenti-shRNA-ephrinB2: 2.33 ± 7.44 days, n = 9 mice, p = 0.099, unpaired t-test) (Fig 4j). Lastly, given that we targeted the location of the critically-important pool of PhMNs with our virus injections, we determined whether ephrinB2 knockdown specifically within the cervical ventral horn extended animal survival, as determined by the righting reflex ^29,31^. Compared to lenti-GFP control, lenti-shRNA had no effect on the age of disease endstage as determined by Kaplan-Meier analysis (Lenti-GFP: 132.0 days; Lenti-shRNA-ephrinB2 136.5 days, chi square: 0.24, p = 0.63, Gehan-Breslow-Wilcoxon test; n = 8-10 mice per group) (Fig 4k).

**Figure 4:**
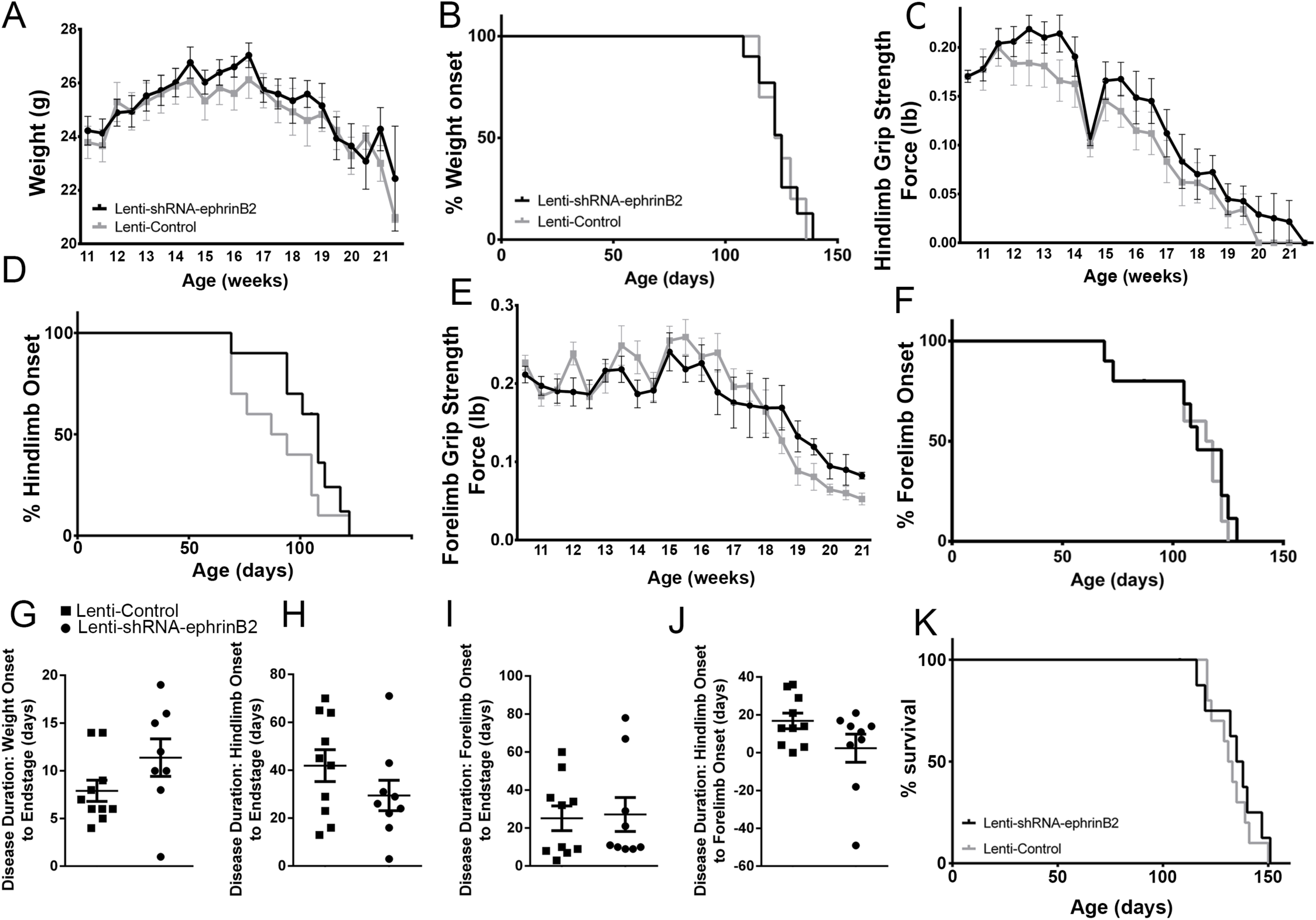
Knockdown of ephrinB2 in the cervical spinal cord ventral horn did not extend survival or delay onset of disease in SOD1^G93A^ mice. Biweekly weights of Lenti-shRNA-ephrinB2 and Lenti-Control mice were recorded until endpoint sacrifice (**a**), and disease onset was measured for each animal when there was a 10% drop in body weight (**b**). Biweekly individual hindlimb grip strengths were assessed for each animal (**c**), and disease onset was recorded when the animal had a 10% decline in hindlimb grip strength (**d**). Each animal was also tested for forelimb grip strength (**e**), and disease onset was recorded when the animal had a 10% decline in forelimb grip strength (**f**). Weights, forelimb grip strength and hindlimb grip strength were taken biweekly starting one week prior to injection of Lenti-shRNA-ephrinB2 or Lenti-GFP control, and all force measurements plotted were the average force (lb) of all animals combined in each group. Disease duration was determined by time from weight onset to endstage (**g**), hindlimb onset to endstage (**h**), and forelimb onset to endstage for each animal (**i**). Disease duration was also measured from the time of forelimb onset to time of hindlimb onset (**j**). Survival was measured as the day each animal reached endstage, which was determined by the righting reflex (**k**). Analyses in all panels: n = 8-10 mice per genotype and per time point; 4-5 females and 4-5 males per condition.

### Preservation of PhMN innervation of the diaphragm

We next quantified morphological innervation changes at the diaphragm NMJ, as this synapse is critical for functional PhMN-diaphragm circuit connectivity. We labeled phrenic motor axons and their terminals for neurofilament (using SMI-312R antibody) and synaptic vesicle protein 2 (SV2), respectively, and we labeled nicotinic acetylcholine receptors with Alexa555-conjugated alpha-bungarotoxin ^27,30^. Using confocal imaging of individual NMJs, we quantified the percentage of intact (Fig 5a), partially-denervated (Fig 5b) and completely-denervated (Fig 5c) NMJs in the diaphragm ^37–39^. Though we assessed NMJ morphology only in SOD1^G93A^ mice in this work, our previous findings show that all diaphragm NMJs in non-diseased wild-type mice are completely intact ^28,30,40,41^. In the current study, we find extensive denervation at a large portion of NMJs across the diaphragm at 117 days of age in SOD1^G93A^ mice. Compared to control-treated animals (Fig 5d), the lenti-shRNA group (Fig 5e) showed a significant increase in the percentage of fully-innervated NMJs (Fig 5f) and a significant decrease in percentage of completely-denervated junctions (Fig 5g), demonstrating that lenti-shRNA treatment preserved PhMN innervation of the diaphragm (innervated: Lenti-GFP: 27.0 ± 2.5 % of total NMJs, n = 4 mice; Lenti-shRNA: 53.2 ± 8.5, n = 4; t-test, p = 0.04) (denervated: Lenti-GFP: 21.6 ± 1.6 % of total NMJs, n = 4 mice; Lenti-shRNA: 8.0 ± 3.7, n = 4 mice; t-test, p = 0.03). We also found a trend toward a decrease in the percentage of partially-denervated NMJs in lenti-shRNA animals versus control (Fig 5h), though the difference was not significant (partially-denervated: Lenti-GFP: 42.1 ± 0.7 % of total NMJs, n = 4 mice; Lenti-shRNA: 29.6 ± 6.2; n = 4 mice, t-test, p = 0.11). Our NMJ analyses suggest that preservation of diaphragm innervation by PhMNs with focally- delivered lenti-shRNA-ephrinB2 resulted in a maintenance of diaphragm function. The increased cervical MN survival in the Lenti-shRNA-ephrinB2 group coincided with enhanced preservation of diaphragm NMJ innervation, suggesting that the ephrinB2 knockdown-mediated effects on NMJ innervation and CMAP amplitudes were due at least in part to protection of PhMNs centrally within the cervical spinal cord.

**Figure 5:**
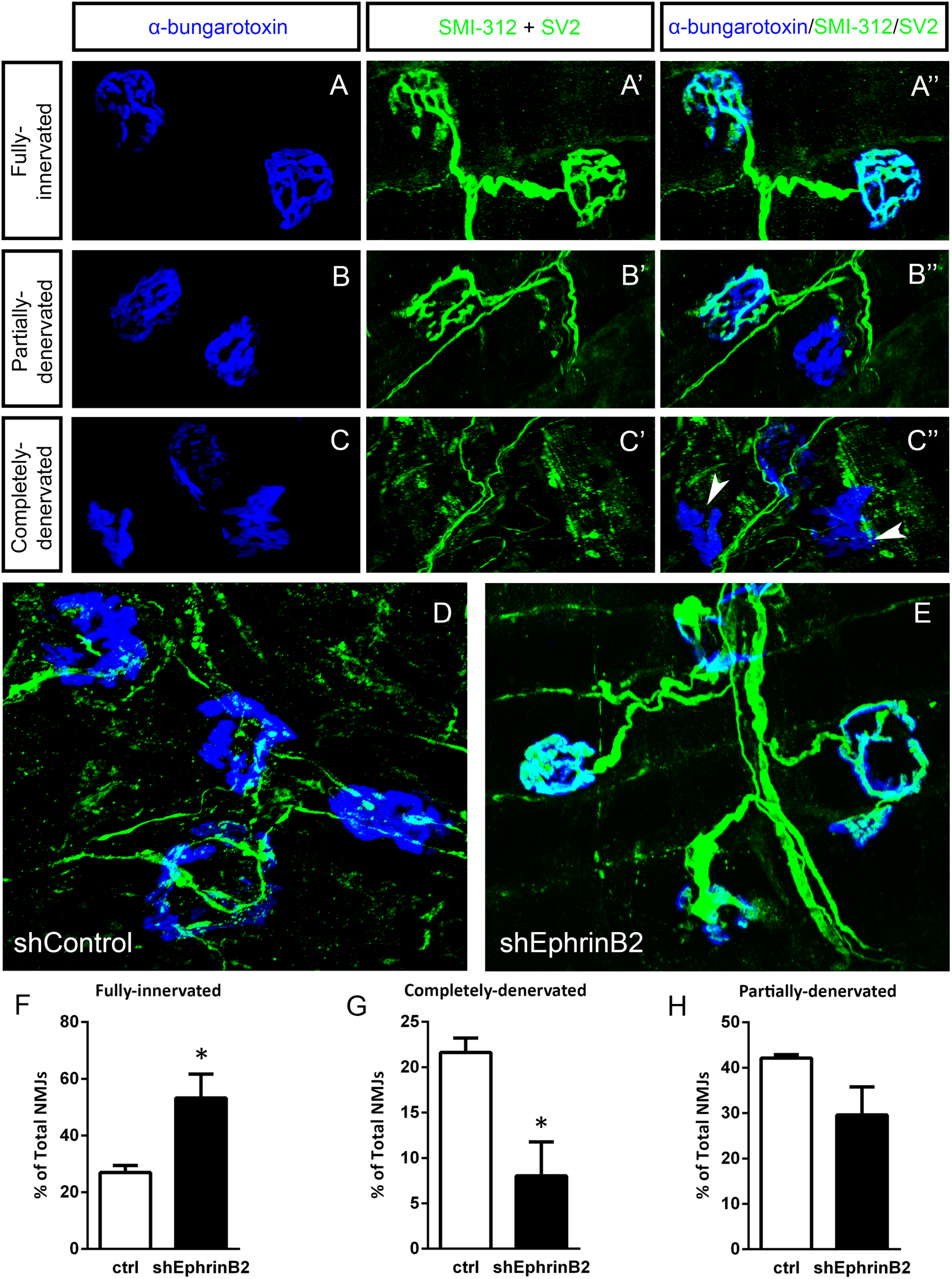
EphrinB2 knockdown preserved morphological innervation of the diaphragm in SOD1^G93A^ mice. Diaphragm muscles were labeled with SMI-312 (green), SV2, (green) and alpha-Bungarotoxin (blue). Representative images of fully-innervated (**a**), partially-denervated (**b**) and completely-denervated (**c**: arrowheads denote completely-denervated NMJs) NMJs are shown; scale bar: 20 µm. Compared to SOD1^G93A^ mice treated with Lenti-GFP (**d**), animals injected with Lenti- shRNA-ephrinB2 (**e**) showed greater preservation of PhMN innervation of the diaphragm NMJ. Quantification revealed a significant increase in the percentage of fully-innervated NMJs (**f**) and a decrease in the percentage of completely-denervated NMJs (**g**) in the Lenti-shRNA-ephrinB2 group compared to Lenti-GFP controls. The percentage of partially-denervated NMJs was not statistically different between the two groups (**h**). Analyses in all panels: n = 4 mice per condition; 2 females and 2 males per condition.

### EphrinB2 upregulation in human ALS spinal cord

We also performed immunoblotting analysis on postmortem samples from human ALS donors with an SOD1 mutation (n=3 donors) and matching non-diseased human samples (n=3 donors). In the lumbar enlargement, there was a large increase in ephrinB2 protein expression in the SOD1 mutation ALS samples compared to the non-diseased controls (Fig 6) (non-ALS lumbar: 3.8 ± 1.1 a.u.; ALS lumbar: 17.8 ± 18.2; ALS cortex: 2.9 ± 0.25; p = 0.523, ANOVA). There was some donor-to-donor variability; while all of the non-ALS control samples showed similarly lower levels of ephrinB2 protein expression in the lumbar spinal cord, dramatic ephrinB2 upregulation in the SOD1 mutation samples was observed with only two of the three ALS donors. The absence of ephrinB2 upregulation in the one ALS sample may be related to the anatomical progression of disease in this particular donor. To this point, we also performed GFAP immunoblotting on the same lumbar spinal cord samples and found signficantly higher GFAP protein levels in the two samples with increased ephrinB2 expression (Fig 6). As the level of GFAP expression is often used as an indicator of disease progression at a particular anatomical region, this finding suggests that ephrinB2 upregulation may have occurred selectively at locations in the CNS where disease processes were already occurring by the time of death. Lastly, we did not observe increased ephrinB2 expression in a disease unaffected region in these same three ALS donor samples, as ephrinB2 protein levels were not elevated in frontal cortex (Fig 6).

**Figure 6:**
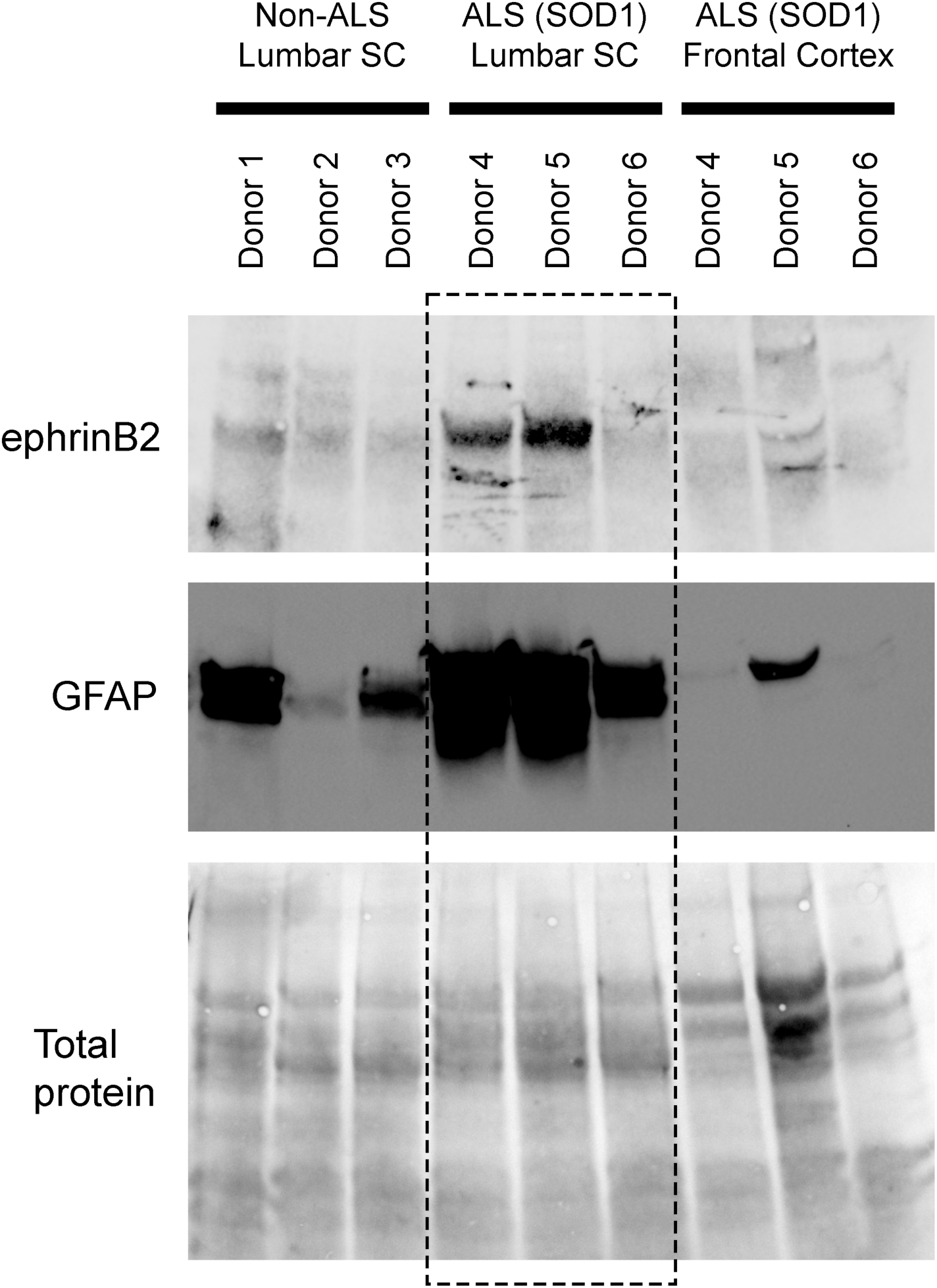
EphrinB2 upregulation in human spinal cord from ALS donors with an SOD1 mutation. Immunoblotting analysis on postmortem samples from human ALS donors with an SOD1 mutation and also non-diseased human samples. In the lumbar enlargement, there was a large increase in ephrinB2 protein expression in the SOD1 mutation ALS samples compared to the non-diseased controls (top blot). GFAP immunoblotting on the same lumbar spinal cord samples shows a robust increase in GFAP protein levels in the two samples with increased ephrinB2 expresison (middle blot). There was not increased ephrinB2 expression in a disease unaffected region in these same three ALS donor samples, as ephrinB2 protein levels were not elevated in the frontal cortex (top blot). Immunoblot for total protein (bottom blot). Demographic information: **Donor 1** – death at 67 years; male; non-ALS; **Donor 2** – death at 70 years; male; non-ALS; **Donor 3** – death at 70 years; female; non-ALS; **Donor 4** – death at 42 years; female; SOD1-D102H mutation; absence of C9orf72 repeat expansion; **Donor 5** – death at 55 years; male; SOD1-A4V mutation; absence of C9orf72 repeat expansion; **Donor 6** – death at 58 years; male; SOD1-V87A mutation; absence of C9orf72 repeat expansion.

## DISCUSSION

We have shown that ephrinB2 is expressed predominantly by ventral horn astrocytes and that ephrinB2 up-regulation coincided with progression of MN loss and overall disease phenotype in the SOD1^G93A^ mouse model of ALS. Furthermore, we found that reducing ephrinB2 expression in ventral horn astrocytes of the SOD1^G93A^ mouse cervical spinal cord maintained diaphragmatic respiratory function by protecting cervical spinal cord MNs and preserving PhMN innervation of the diaphragm. Despite the significant impact that ephrinB2 knockdown had on functional diaphragm innervation, we did not observe effects on limb motor function, disease onset, phenotypic progression of disease post-onset, or overall animal survival. Measures such as disease duration and animal survival also depend on other MN populations such as those present in lumbar spinal cord, brainstem and motor cortex, while our targeted strategy only addresses cervical MN loss (and more specifically those MNs located only at C3-C5). That being said, PhMN preservation plays a critical role in both SOD1^G93A^ models and human disease ^35^, and we have previously shown that focal protection of cervical enlargement MNs using glial progenitor transplantation does extend overall disease phenotype in SOD1^G93A^ rodents ^25^. Nevertheless, in future work aimed at addressing more translational considerations, we can extend this approach to delivery strategies such as intrathecal injection to target ephrinB2 throughout the spinal cord neuraxis. These data with focal ephrinB2 knockdown demonstrate the important role played by ephrinB2 in MN health and muscle function in mutant SOD1-associated ALS pathogenesis and suggest that ephrinB2 is a promosing target for further investigation.

Other than in a relatively small number of studies ^16–20,32,42,43^, the role of Eph-ephrin signaling in ALS has not been extensively examined. Our findings suggest that astrocyte ephrinB2 may play a non-cell autonomous role in ALS, in particular in mutant SOD1-associated disease. A substantial body of work has demonstrated that astrocytes are involved in ALS pathogenesis ^7,8^, both via loss of critically-important functions such as extracellular glutamate uptake ^44^ and toxic gain-of-function such as altered transforming growth factor β (TGFβ) signaling ^45^ and increased production of reactive oxygen species (ROS) ^46^. The dramatic increase of ephrinB2 expression observed in ventral horn astrocytes may represent an additional toxic property.

What is the mechanism by which astrocyte ephrinB2 contributes to MN pathology in ALS? EphA4 receptor expression and signaling capacity correlate with the degree of human ALS disease severity, and EphA4 significantly contributes to MN degeneration in several animal models of ALS ^16^. EphA4 receptor can be activated by ephrinA and ephrinB ligands including ephrinB2 ^47^, suggesting that astocyte ephrinB2 serves as a ligand mediating pathogenic actions in ALS. Consistent with this model, ephrinB2 is upregulated in the SOD1^G93A^ mouse model and reduction of ephrinB2 expression increases MN surivival near the site of ephrinB2 knockdown. Going forward, we could explore this potential interaction in vivo and in astrocyte-MN co-cultures using approaches such cell type-specific knockout of ephrinB2 and EphA4 expression, preventing ephrinB2-EphA4 binding, and EphA4 receptor kinase activity indicators.

Previous findings suggest that EphA-ephrinB signaling may contribute to ALS pathogenesis. Initial results showed that EphA4 plays a significant role in both ALS animal models and in the human ALS population ^16^. Subsequent work in mouse models ^18–20^ showed that genetic reduction of EphA4 in SOD1^G93A^ mice or intracerebroventricular delivery of an antisense oligonucleotide directed against EphA4 in both SOD1^G93A^ and PFN1^G118V^ mouse models of ALS did not impact disease measures. In contrast, manipulations that target EphA4 signaling such as administration of an EphA4 agonist to mutant SOD1 mouse increased both disease duration and animal survival ^48^. In addition, delivery of soluble EphA4-Fc that blocks ligand binding to EphA4 results in partial preservation of motor function in SOD1^G93A^ mice ^20^. Importantly, these inhibitors act via blocking the Eph-ephrin interaction or disrupt bidirectional Eph-ephrin signaling. Thus, these data are consistent with a model where reverse or bidirectional EphA-ephrinB signaling via the interaction of EphA4 with ephrinB2 may be involved in ALS. Supporting this notion, genetic knockdown of ephrinA5 (an EphA4 ligand) in the SOD1^G93A^ mouse model accelerates disease progression and hastens animal death ^17^, which may be explained by an enhancement of the EphA4-ephrinB2 interaction in the absence of ephrinA5. While other agents are also being developed to manipulate binding of ephrins with EphA4 for ALS therapeutics ^49,50^, our results suggest that directly targeting ephrinB2 is a promising strategy to modulate both Eph-ephrin signaling and astrocyte-MN interactions in ALS. Unlike the effects of knocking down EphA4 expression in ALS animal models, we observe significant MN protection, maintenance of NMJ innervation and preservation of diaphragm muscle function following ephrinB2 reduction.

In addition to EphA4, ephrinB2 could signal via another Eph-family protein in ALS. Consistent with this model, ephrin-B upregulation in chronic pain models results in increased NMDA receptor function and pathological synaptic plasticity via interaction with EphBs ^51^. EphB’s are centrally involved in regulating subcellular localization of ionotropic glutamate receptor subunits to excitatory synapses ^52,53^, raising the intriguing possibility that enhanced ephrinB2-EphB2 signaling results in increased glutamate receptor activation in MNs and consequently may be contributing to excitotoxicity that plays a well-known role in ALS. Thus, ephrinB2 upregulation provides a number of potential avenues for aberrant circuit plasticity that could enhance neuronal damage and contribute to ALS pathogenesis.

In this study, we did not examine Eph receptor expression in the PhMN pool, but only focused on ephrinB2 in the surrounding astrocytes. Nevertheless, there are several pieces of evidence to suggest that PhMNs do express a variety of Eph receptors, which are possible candidates through which astrocyte ephrinB2 exerts its actions on PhMN health. EphrinB2 binds and activates EphBs, as well as EphAs such as EphA4. Importantly, previous studies have linked expression of EphA4 in MNs to the rate of ALS progression ^16^. Consistent with these studies, single-nucleus RNAseq on mouse cervical spinal cord shows that alpha MNs of the cervical spinal cord express various EphA and EphB receptors (http://spinalcordatlas.org/) ^54,55^. In addition, this dataset identifies a PhMN-specific marker (ErbB4); by specifically looking at the expression profile of only the ErbB4-expressing cervical alpha MNs, the data reveal that PhMNs express a number of EphA’s and EphB’s, including EphA4. To validate the expression specifically of EphA4, we find via IHC analysis that large ventral horn neurons are positive for phosphorylated EphA4 (a marker of activated EphA4) in C3-C5 spinal cord sections from the SOD1^G93A^ mice injected with shRNA-ephrinB2 vector or control vector (data not shown).

These cervical spinal cord levels include MN pools in addition to just the PhMNs; therefore, this result by itself does not conclusively show that PhMNs at this location express EphA4, though they likely do since we find EphA4 expression in most large neuron cell bodies in C3-C5. Collectively, these RNAseq and IHC data show that PhMNs express a number of Eph receptors, which could be involved in direct cell-cell signaling with surrounding ephrinB2-expressing astrocytes.

EphrinB2 contribution to ALS is likely an astrocyte-mediated phenomenon given the pronounced upregulation that occurs almost entirely in ventral horn astrocytes. Nevertheless, our shRNA vector does not exclusively target only astrocytes. While the majority of transduced cells are astrocytes, we did not identify the lineage of a portion of the transduced cells, which could consist of cell types such as microglia ^56^, endothelial cells ^57^ and others, some of which have been linked to ALS pathogenesis. It is therefore possible that the effects of ephrinB2 knockdown may not only be due to effects on astrocytes. However, we observed (1) dramatic ephrinB2 upregulation in the ventral horn of SOD1^G93A^ mice that appears to be astrocyte-specific, (2) transduction predominately of astrocytes with our vector, and (3) significant reduction of ephrinB2 expression almost entirely in astrocytes in shRNA-treated animals. These data suggest that the effect of ephrinB2-shRNA treatment was primarily due to changes in astrocytes and that astrocyte expression is the likely mechanism underlying ephrinB2’s action in mutant SOD1-associated ALS. In future work, could address this point by employing alternative vector-based strategies or cell type-specific knockout mouse models to selectively target astrocytes alone. In the present study, we used a viral vector with the U6 type-III RNA polymerase III promotor given its utility for persistently expressing high levels of non-coding RNA transcripts such as shRNA ^58^. Despite being a promotor that is not lineage-specific, we found that the majority of tranduced cells were GFAP-positive, though this transduction was still not completely limited to astrocytes. In our previous work, we instead used the Gfa2 promotor to direct highly astrocyte-specific transduction in the adult rodent spinal cord ^40,59–61^, which is an example of an approach moving forward to achieve astrocyte ephrinB2 knockdown in ALS animal models.

We previously showed that there is a modest increase in astrocyte number in ventral horn of the SOD1^G93A^ mouse model at time points following phenotypic disease onset ^62^. It is therefore possible that the increased ephrinB2 expression observed across the ventral horn in SOD1^G93A^ animals in the present study was due to this increased astrocyte number. However, this is unlikely to be the case, as astrocytes (and all other spinal cord cell types) in wild-type mice and in SOD1^G93A^ mice prior to disease onset express very low levels of ephrinB2. Throughout disease course in these SOD1^G93A^ mice, ephrinB2 level in individual astrocytes dramatically increases (including across most or all astrocytes), suggesting that the total increase in ephrinB2 expression across the ventral horn was not due to just this small increase in astrocyte numbers but was instead due to the dramatically elevated ephrinB2 expression observed across the astrocyte population.

In this study, we have shown that ephrinB2 protein expression is significanty increased in the lumbar spinal cord of postmortem samples from human ALS donors with an SOD1 mutation compared to non-diseased human samples, suggesting that this disease mechanism is also relevant to the human condition and is not restricted to only the mutant SOD1 mouse model. ALS is heterogeneous both with respect to its genetic basis and its clinical disease course (e.g., age and site of onset; severity/progression) ^23^. The majority of patients have sporadic disease that is not linked to a known heritable genetic cause, while the remaining cases are linked to a known familial genetic mutation.

Furthermore, these familial cases are associated with mutations in a number of different genes. In addition, patients (even with mutations in the same gene) show variability in their clinical disease manifestation such as the rate of disease progression depending on, for example, the specific SOD1 mutation ^63,64^. However, the mechanisms underlying this heterogeneity in human disease progression are not understood. An important consideration is whether ephrinB2’s function is specific to mutant SOD1-mediated disease or extends to more subtypes of ALS, including other disease-associated genes and sporadic ALS. To address whether ephrinB2 is a general modifier of ALS, future studies should focus on post-mortem tissue samples and pluripotent stem cell-derived astrocytes and MNs derived from patients with various subtypes of the disease, as well as on animal models involving other ALS-associated genes. Previous work in ALS8-linked ALS (a form of familial ALS associated with the VAMP-associated protein B gene) suggests the possible relevance of Eph/ephrin biology to ALS pathogenesis ^32^. In addition, EphA4 knockdown can protect against the axonal damage response elicited by expression of ALS-linked mutant TDP-43 ^16^. These data suggest that altered Eph-ephrin signaling may not be limited to only mutant SOD1-assocated ALS, though more extensive investigation is necessary to support this idea.

Respiratory function involves the contribution of a number of other muscle groups, and these muscles are innervated by various lower MN pools located across a relatively-large expanse of the CNS neuraxis. While CMAP recording is a powerful assay of functional innervation of diaphragm muscle by phrenic motor axons, it does not directly measure overall respiratory function. There are assays to test outcomes such as ventilatory behavior and gas exchange (e.g., whole-body plethysmography, blood gas measurements, etc.) ^26^. As we focally targeted our ephrinB2 knockdown to only a small area (the phrenic nucleus), we would not expect an effect on these other functional assays, which is why we restricted our functional testing to CMAP recording to specifically study the effects of ephrinB2 knockdown on the PhMN pool.

Interesingly, we observed relatively robust effects of focal ephrinB2 knockdown in the cervical enlargement on functional diaphragm innervation, but did not similarly find effects on forelimb motor function using the forelimb grip strength assay, despite forelimb-innervating MN pools also residing in the cervical spinal cord. However, this functional assay is impacted more by distal forelimb muscle groups controlled by MN pools located at more caudal locations of the spinal cord (i.e. low cervical and high thoracic), likely explaining the lack of effect on grip strength. The localized – yet robust – effects of ephrinB2 knockdown are consistent with the model that ephrinB2 is a target worth further exploration and validation.

In summary, we found astrocyte-specific upregulation of ephrinB2 expression in the ALS spinal cord, and we demonstrated that knocking down ephrinB2 in the ventral horn in an anatomically-targeted manner significantly preserved diaphragmatic respiratory neural circuitry in SOD1^G93A^ mice.

Importantly, ephrinB2 knockdown exerted significant protective effects on the centrally-important population of respiratory PhMNs, which translated to maintenance of diaphragm function *in vivo*. We also report significantly increased ephrinB2 expression in the disease affected spinal cord of mutant SOD1 human ALS samples. In conclusion, our findings suggest that astrocyte ephrinB2 upregulation is both a signaling mechanism underlying astrocyte pathogenicity in mutant SOD1-associated ALS and a promising therapeutic target.

## MATERIALS AND METHODS

### Animal model

Female and male transgenic SOD1^G93A^ mice (C57BL/6J congenic line: B6.Cg-Tg(SOD1*G93A)1Gur/J and B6SJL-Tg(SOD1*G93A)1Gur/J) were used in all experiments. All procedures were carried out in compliance with the National Institutes of Health (NIH) Guide for the Care and Use of Laboratory Animals and the ARRIVE (*Animal Research: Reporting of In Vivo Experiments*) guidelines. Experimental procedures were approved by Thomas Jefferson University Institutional Animal Care and Use Committee (IACUC). All animals were housed in a temperature-, humidity-, and light-controlled animal facility and were provided with food and water *ad libitum*.

### Endstage care

Due to the progression of muscle paralysis, animals were given access to softened food and were checked daily for overall health once the animals reached phenotypic onset of disease. We determined the onset for each animal by assessing total weight, hindpaw grip strength and forepaw grip strength (described below) ^25,40^. Animals were considered to have reached onset when there was a 10% loss in total body weight or a 10% loss in either forelimb or hindlimb grip strength. To determine the endstage for each animal, we used the “righting reflex” method. We placed animals on their left and right sides; if a mouse could not right itself after 30 seconds on both sides, it was euthanized with an overdose of ketamine/xylazine.

### Viral vectors

Vectors used were VSVG.HIV.SIN.cPPT.U6.SbRmEphrinB2.4.CMV.EGFP and VSVG.HIV.SIN.cPPT.U6.Empty.CMV.EGFP. Lenti-shRNA-ephrinB2 or Lenti-Control constructs were driven by the U6 promoter, and EGFP expression was driven by the cytomegalovirus (CMV) promoter ^24^. shRNA sequence: ephrin-B2 shRNA: 5-GCAGACAGATGCACAATTA-3. Forward and reverse oligonucleotides were synthesized (Integrated DNA Technologies) and generated a dsDNA insert consisting of forward and reverse complement RNAi sequences separated by a hairpin region and flanked by restriction site overhangs. We used 1.9 × 10^^10^ for intraspinal injections (described below).

### Intraspinal injection

For the intraspinal injections ^65–67^, mice were first anesthetized with 1% isoflurane in oxygen, and the dorsal surface of the skin was shaved and cleaned with 70% ethanol. A half-inch incision was made on the dorsal skin starting at the base of the skull, and the underlying muscle layers were separated with a sterile surgical blade along the midline between the spinous processes of C2 and T1 to expose the cervical laminae. Paravertebral muscles overlying C3-C5 were removed using rongeurs, followed by bilateral laminectomies of the vertebrae over the C3-C5 spinal cord. A 33-gauge (G) needle on a Hamilton microsyringe (Hamilton, Reno, Nevada) was lowered 0.8 mm ventral from the dorsal surface just medial to the entry of the dorsal rootlets at C3, C4 and C5. After inserting the needle into the ventral horn, we waited three minutes before injecting the viral constructs. 2 uL of Lenti-shRNA-ephrinB2 or Lenti-Control virus were delivered to the spinal cord over 5 minutes, controlled by an UltraMicroPump and Micro4 Microsyringe Pump Controller (World Precision Instruments, Sarasota, Florida). After injection, the needle was left in place for 3 minutes before being slowly removed. Following intraspinal injection, dorsal muscle layers were sutured with 4-0 silk sutures (Covidien, Minneapolis, Minnesota) and the skin was closed with surgical staples (Braintree Scientific, Braintree, Massachusetts). The surface of the skin was treated with a topical iodine solution. Immediately following the procedure, mice were given 1 mL of Lactated Ringer’s solution (Hospira, San Jose, California) and cefazolin (6 mg) (Hospira, San Jose, California) via subcutaneous injections. Mice were placed in a clean cage on a surgical heating pad set to 37° C (Gaymar, Orchard Park, New York). At 12 and 24 hours after surgery, each animal was given an additional dose of buprenorphine hydrochloride (0.05mg/kg) and monitored for pain/distress. Mice were 60 days old at the time of virus injection.

### Weight and grip-strength test

Weights were measured for each animal biweekly prior to forelimb and hindlimb testing. Forelimb and hindlimb grip strengths were determined using a “Grip Strength Meter” (DFIS-2 Series Digital Force Gauge; Columbus Instruments, OH) ^29,67^. Grip strength was measured by allowing the animals to tightly grasp a force gauge bar using both forepaws or both hindpaws, and then pulling the mice away from the gauge until both limbs released the bar. The force measurements were recorded in three trials, and the averages were used in analyses. Grip strengths were recorded biweekly starting one week prior to initial injection.

### Compound Muscle Action Potential (CMAP) recordings

At 117 days of age, mice were anesthetized with isoflurane (Piramal Healthcare, Bethlehem, Pennsylvania) at a concentration of 1.0-1.5% in oxygen. Animals were placed supine, and the abdomen was shaved and cleaned with 70% ethanol. Phrenic nerve conduction studies were performed with stimulation of the phrenic nerve via needle electrodes trans-cutaneously inserted into the neck region in proximity to the passage of the phrenic nerve ^68,69^. A reference electrode was placed on the shaved surface of the right costal region. Phrenic nerve was stimulated with a single burst at 6mV (amplitude) for a 0.5 millisecond duration. Each animal was stimulated between 10-20 times to ensure reproducibility, and recordings were averaged for analysis. ADI Powerlab8/30stimulator and BioAMPamplifier (ADInstruments, Colorado Springs, CO) were used for both stimulation and recording, and Scope 3.5.6 software (ADInstruments, Colorado Springs, CO; RRID: SCR_001620) was used for subsequent data analysis. Following recordings, animals were immediately euthanized, and tissue was collected (as described below).

### Diaphragm dissection

Animals were euthanized by an intraperitoneal injection of ketamine/xylazine diluted in sterile saline and then placed in a supine position. A laparotomy was performed to expose the inferior surface of the diaphragm. The diaphragm was then excised using spring scissors (Fine Science Tools, Foster City, California), stretched flat and pinned down on silicon-coated 10 cm dishes, and washed with PBS (Gibco, Pittsburgh, Pennsylvania). Diaphragms were then fixed for 20 minutes in 4% paraformaldehyde (Electron Microscopy Sciences, Hatfield, Pennsylvania). After washing in PBS, superficial fascia was carefully removed from the surface of the diaphragm with Dumont #5 Forceps (Fine Science Tools, Foster City, California). Diaphragms were then stained for NMJ markers (described below).

### Diaphragm whole-mount histology

Fresh diaphragm muscle was dissected from each animal for whole-mount immunohistochemistry, as described above ^68,70^. Diaphragms were rinsed in PBS and then incubated in 0.1 M glycine for 30 minutes. Following glycine incubation, α-bungarotoxin conjugated to Alexa Fluor 555 at 1:200 (Life Technologies, Waltham, Massachusetts) was used to label post-synaptic nicotinic acetylcholine receptors. Ice-cold methanol was then added to the diaphragms for 5 minutes, and then diaphragms were blocked for 1 hour at room temperature in a solution of 2% bovine serum albumin and 0.2% Triton X-100 diluted in PBS (this solution was used for both primary and secondary antibody dilutions). Primary antibodies were added overnight at 4° C: pre-synaptic vesicle marker anti-SV2 at 1:10 (Developmental Studies Hybridoma Bank, Iowa City, Iowa; RRID: AB_2315387); neurofilament marker anti-SMI-312 at 1:1000 (Covance, Greenfield, Indiana; RRID: AB_2314906). The diaphragms were then washed and secondary antibody solution was added for 1 hour at room temperature: FITC anti-mouse IgG secondary (Jackson ImmunoResearch Laboratories, West Grove, PA; 1:100). Diaphragms were mounted with Vectashield mounting medium (Vector Laboratories, Burlingame, California), coverslips were added, and slides were stored at −20°C.

### Neuromuscular junction (NMJ) analysis

At 117 days of age, labeled muscles were analyzed for the percentage of NMJs that were intact, partially-denervated or completely denervated ^30,38^. Whole-mounted diaphragms were imaged on a FV1000 confocal microscope (Olympus, Center Valley, Pennsylvania; RRID: SCR_014215). We conducted NMJ analysis on the right hemi-diaphragm.

### Spinal cord and brain dissection

Animals were euthanized by an intraperitoneal injection of ketamine/xylazine diluted in sterile saline (as described above). Following diaphragm removal (described below), the animal was exsanguinated by cutting the right atrium and transcardially perfused with 0.9% saline solution (Fisher Scientific, Pittsburgh, Pennsylvania) then 4% paraformaldehyde (Electron Microscopy Sciences, Hatfield, Pennsylvania) to fix the tissue. Following perfusion, the spinal cord and brain were excised with rongeurs (Fine Science Tools, Foster City, California) and kept in a 4% paraformaldehyde solution overnight at 4° C, washed with 0.1 M Phosphate Buffer (Sodium Phosphate Dibasic Heptahydrate (Sigma-Aldrich, St. Louis, Missouri) and Sodium Monobasic Monohydrate (Sigma-Aldrich, St. Louis, Missouri)), and placed in 30% sucrose (Sigma-Aldrich, St. Louis, Missouri). A second group of animals was not perfused with 4% paraformaldehyde, and brain and spinal cord tissue were collected unfixed. Both fixed and unfixed samples were placed into an embedding mold (Polysciences Inc, Warrington, Pennsylvania) and covered with tissue freezing medium (General Data, Cincinnati, Ohio). Samples were then flash frozen in 2-methylbutane (Fisher Scientific, Pittsburgh, Pennsylvania) chilled in dry ice. Tissue was sectioned at 30 µm on a cryostat (Thermo Scientific, Philadelphia, Pennsylvania), placed on glass microscope slides (Fisher Scientific, Pittsburgh, Pennsylvania), and dried overnight at room temperature before freezing the samples at −20° C for long term storage.

### Spinal cord histology/cresyl violet staining

Spinal cord tissue section slides were dried at room temperature for 2 hours. Following drying, slides were rehydrated in 3-minute baths of xylene, 100% ethanol, 95% ethanol, 70% ethanol and dH2O. To stain the tissue, slides were placed in an Eriochrome solution (0.16% Eriochrome Cyanine, 0.4% Sulfuric Acid, 0.4% Ferric Chloride in dH2O) for 14 minutes, washed with tap water, placed in a developing solution (0.3% ammonium hydroxide in dH2O) for 5 minutes, washed with dH2O, and then placed into a cresyl violet solution (0.4% cresyl violet, 6% 1M sodium acetate, 34% 1M acetic acid) for 18 minutes. After staining, slides were dehydrated by being placed in baths of dH2O, 70% ethanol, 95% ethanol, 100% ethanol and xylene. Slides were mounted with poly-mount xylene (Polysciences, Warrington, Pennsylvania), and cover slips were added. Slides were then kept at room temperature for storage and analysis.

### Immunohistochemistry

Prior to immunostaining, tissue sections were dried for 1 hour at room temperature. Antigen retrieval was performed using R&D Systems Protocol (R&D Systems, Minneapolis, Minnesota). Immediately after antigen retrieval, a hydrophobic pen (Newcomer Supply, Middleton, Wisconsin) was used to surround the tissue sections. Slides were blocked/permeabilized for 1 hour at room temperature with a solution of 5% Normal Horse Serum (Vector Laboratories, Burlingame, California), 0.2% Triton X-100 (Amresco, Solon, Ohio), diluted in PBS (primary and secondary antibodies were diluted in this solution as well). Slides were then treated with primary antibody overnight at 4° C with the following antibodies: neuronal marker anti-NeuN at 1:200 (EMD-Millipore, Temecula, California; AB_2298772); astrocyte marker anti-GFAP at 1:400 (Dako, Carpinteria, California; RRID: AB_10013482); oligodendrocyte lineage marker anti-Olig-2 at 1:200 (EMD-Millipore, Temecula, California; RRID: AB_2299035); anti-ephrinB2 at 1:50 (R&D Systems, Minneapolis, Minnesota, RRID: AB_2261967); anti-ephA4 at 1:100 (R&D Systems, Minneapolis, Minnesota RRID: AB_2099371); and anti-GFP at 1:500 (Aves Labs, Davis, California, RRID: AB_10000240). On the following morning, samples were washed 3x in PBS, and secondary antibody solutions were added for 1 hour at room temperature: donkey anti-rabbit IgG H&L (Alexa Fluor 647) at 1:200 (Abcam, Cambridge, Massachusetts); donkey anti-mouse IgG H&L (Alexa Fluor 488) at 1:200 (Abcam, Cambridge, Massachusetts); Rhodamine (TRITC) AffiniPure donkey anti-goat IgG (H+L) at 1:200 (Jackson ImmunoResearch, West Grove, Pennsylvania). Following secondary antibody treatment, samples were washed in PBS and 2 drops of FluorSave reagent (Calbiochem, San Diego, California) were added to tissue sections, then slides were coverslipped (Fisher Scientific, Pittsburgh, Pennsylvania). Slides were stored at 4° C.

### Viral vector transduction quantification

SOD1^G93A^ mouse cervical spinal cord tissue at disease endstage was immunostained with anti-GFP and either anti-GFAP, anti-NeuN or anti-Olig2 (described above). We quantified the percentage of double-labeled GFP+/GFAP+, GFP+/NeuN+ or GFP+/Olig2+ cells versus the total number of GFP+ cells in the ventral horn. The cell lineage of lenti- viral transduction was plotted as a percentage of the total GFP+ cells.

### Motor neuron counts

At 117 days of age, 30 µm mouse cervical spinal cord tissue sections were stained with cresyl violet (as described above) to determine the total number of motor neurons.

Images were acquired using a 10x objective on a Zeiss Axio M2 Imager (Carl Zeiss Inc., Thornwood, New York), and analyzed with ImageJ/Fiji software (RRID: SCR_003070). The area (converted into pixels) of each ventral horn was outlined separately starting from the central canal and tracing laterally and ventrally to encompass the right and left ventral horns for each spinal cord section.

Within the area of each ventral horn, neurons were traced and somal area was assessed. We considered a motor neuron as any neuron within the ventral horn greater than 20 µm in somal diameter and with an identifiable nucleolus ^40^. We then assessed total number of motor neurons per area of the ventral horn for both the Lenti-shRNA-ephrinB2 group and the Lenti-control group.

### EphrinB2 quantification

EphrinB2 levels in ventral horn of the cervical spinal cord of endstage SOD1^G93A^ mice intraspinally injected with Lenti-shRNA-ephrinB2 or Lenti-Control were evaluated. In addition, this same analysis was performed on uninjected SOD1^G93A^ mice at 60 days of age, 120 days of age, and at disease endstage, as well as on uninjected wild-type mice at 140 days of age. 30 µm cervical spinal cord sections were immunostained with anti-GFP and anti-ephrinB2 antibodies.

ShRNA-induced knockdown was assessed by quantifying the number of ephinB2+/GFP+ cells for both Lenti-GFP control and Lenti-shRNA-ephrinB2 groups. 4 animals were used for each group, with the number of ephrinB2/GFP+ cells per animal averaged over 3 slides (8 tissue sections each).

### Human postmortem tissue

For analysis of human postmortem tissue, we examined three non-ALS and three ALS donors. Non-diseased samples were obtained from the NIH NeuroBioBank. Age of death for these three non-ALS donors was 67, 70 and 70 years. For the ALS samples, all three donors had an SOD1 mutation (donor 1: D102H mutation; donor 2: A4V; donor 2: V87A) and all did not have a C9orf72 repeat expansion. Two of these SOD1 ALS samples were obtained from Project ALS, and the third sample was obtained from the biorepository of the Jefferson Weinberg ALS Center. These three donors succumbed to ALS at 42 (female), 55 (male) or 58 (male) years of age.

### Immunoblotting of postmortem tissue

100 mg of fresh-frozen human autopsy sample (lumbar spinal cord or frontal cortex) were homogenized in 1% SDS using a Dounce homogenizer. Homogenate was centrifuged at 3000 rpm for 20 minutes at 4 C to remove debris. Clear supernatant was then used to estimate total protein content using the bicinchoninic acid (BCA) assay (Pierce BCA kit #23225; Thermo Fischer Scientific, Waltham, Massachusetts). 30 µg of protein were loaded onto 10% stain-free gel (#4568034; Bio-Rad, Hercules, California). After the run, gels were activated using UV light to crosslink protein and transferred to 0.22 µm nitrocellulose membrane. After transfer, membrane was exposed to chemiluminescence light to image total protein. Membrane was then blocked using 5% fat-free milk in tris-buffered aaline with tween (TBST) for one hour at room temperature. Anti-ephrinB2 antibody (Cat# ab131536, RRID: AB_11156896; Abcam, Cambridge, Massachusetts) at 1:500 dilution in 5% bovine serum albumin in TBST was incubated overnight, followed by three washes with TBST on the shaker for 15 minutes each. Anti-rabbit horseradish peroxidase (HRP) secondary (#NA9340V, Sigma-Aldrich, St. Louis, Missouri) at 1:5000 dilution was prepared in 5% fat-free milk and added to membrane for one hour at room temperature with shaking. Membranes were washed 3x for 15 minutes on a shaker with TBST. Chemiluminescence signal was imaged using super signal west Atto (#38554; Bio-Rad, Hercules, California). The same membrane was used to probe for GFAP using anti-GFAP antibody (#610566; BD Bioscience, Franklin Lakes, New Jersey) at 1:2000 dilution overnight. Membrane was washed 3x the next day with TBST and incubated with anti-mouse HRP (#NXA931V; Sigma-Aldrich, St. Louis, Missouri) at 1:5000 dilution for 1 hour at RT and washed, and then chemiluminescence was imaged as described above. Quantification for ephrinB2 was performed by normalizing to total protein using Bio-rad Image Lab software (RRID:SCR_014210).

### Reagents

We authenticated relevant experimental regents to ensure that they performed similarly across experiments and to validate the resulting data. Whenever we used a new batch of the vector, we verified that the virus performed equivalently from batch-to-batch by confirming in every animal that the vector transduced predominantly GFAP-positive astrocytes and induced similar expression of the GFP reporter for each batch. For Alexa-conjugated α-bungarotoxin and for all antibodies used in the immunohistochemistry studies, we always verified (when receiving a new batch from the manufacturer) that labeling in the spinal cord and/or diaphragm muscle coincided with the established expression pattern of the protein. We have provided Research Resource Identification Initiative (RRID) numbers for all relevant reagents (i.e. antibodies and computer programs) throughout the Materials and Methods section.

### Experimental design and statistical analysis

Before starting the study, mice were randomly assigned to experimental groups, and the different vectors used within a given experiment were randomly distributed across these mice (and within a given surgical day). For all of the phenotypic analyses, we repeated the experiment for both virus groups in two separate cohorts. All surgical procedures and subsequent behavioral, electrophysiological and histological analyses were conducted in a blinded manner. In the Results section, we provide details of exact n’s, group means, standard error of the mean (SEM), statistical tests used and the results of all statistical analyses (including exact p-values, t-values and F-values) for each experiment and for all statistical comparisons. Statistical significance was assessed by analysis of variance (ANOVA) and multiple comparisons *post hoc* test. T-test was used for analysis involving only two conditions. Graphpad Prism 6 (Graphpad Software Inc.; LaJolla, CA; RRID: SCR_002798) was used to calculate all analyses, and *p* ˂ 0.05 was considered significant. While we included both male and female mice, our analyses are under-powered to examine possible sex-specific effects. Given that males make up a larger portion of the human ALS population, it will be important in follow up work to explore the possible sex-specific role of ephrinB2 in ALS pathogenesis.

### Data availability statement

We will make the materials and other resources described in this study available upon reasonable request from academic researchers. In addition, all data associated with this study will be made available in compliance with the FAIR (Findable, Accessible, Interoperable, Reusable) principles. However, we will restrict the information available about the human donors for privacy reasons.

## ACKLOWLEDGEMENTS

This work was supported by the Muscular Dystrophy Association (346986 to A.C.L. and M.B.D.; 628389 to D.T.), the NINDS (R01NS110385 to A.C.L. and M.B.D.; R01NS079702 to A.C.L.; R21NS090912 to D.T.; RF1AG057882 to D.T.; R01NS109150 to P.P.), and the Family Strong for ALS & Farber Family Foundation (P.P., D.T.). Human tissue samples were provided by the NIH NeuroBioBank, Project ALS, and the Jefferson Weinberg ALS Center.

## Author Contributions

Conceptualization: M.B.D., A.C.L.

Methodology: M.W.U., M.B.D., A.C.L.

Software: N.M.H., R.E.C.

Validation: M.W.U.

Formal analysis: M.W.U., M.C.W.

Investigation: M.W.U., B.A.C., N.M.H., S.S.M.

L.S., W.Z., E.V.B., N.T.H., S.J.T., B.G., R.E.C., M.C.W.

Resources: D.T., P.P.

Data curation: M.W.U., M.B.D., A.C.L.

Writing – original draft preparation: M.W.U., A.C.L.

Writing – review & editing: M.W.U., D.T., P.P., M.B.D., A.C.L.

Visualisation: M.W.U., A.C.L.

Supervision: M.B.D., A.C.L.

Project administration: M.B.D., A.C.L.

Funding acquisition: M.B.D., A.C.L.

## ABBREVIATIONS

ALS: amyotrophic lateral sclerosis
C3, 4, 5, etc.: cervical spinal cord level 3, 4, 5, etc.
CMAP: compound muscle action potential
CNS: central nervous system
Eph: erythropoietin-producing human hepatocellular receptor
Ephrin: Eph receptor-interacting protein
G93A: glycine 93 changed to alanine
GFP: green fluorescent protein
GFAP: glial fibrillary acidic protein
MN: motor neuron
NeuN: neuronal nuclear protein
NMJ: neuromuscular junction
Olig2: oligodendrocyte transcription factor 2
PhMN: phrenic motor neuron
shRNA: short hairpin RNA
SMI-312: anti-neurofilament marker
SOD1: superoxide dismutase 1
SV2: synaptic vesicle protein 2
TDP-43: TAR DNA-binding protein 43

## REFERENCES

1. Pekny, M. & Nilsson, M. Astrocyte activation and reactive gliosis. Glia 50, 427–434 (2005).

2. Bruijn, L.I., Miller, T.M. & Cleveland, D.W. Unraveling the mechanisms involved in motor neuron degeneration in ALS. Annu Rev Neurosci 27, 723–749 (2004).

3. Rosen, D.R., et al. Mutations in Cu/Zn superoxide dismutase gene are associated with familial amyotrophic lateral sclerosis. Nature 362, 59–62 (1993).

4. Mackenzie, I.R., et al. Pathological TDP-43 distinguishes sporadic amyotrophic lateral sclerosis from amyotrophic lateral sclerosis with SOD1 mutations. Ann Neurol 61, 427–434 (2007).

5. DeJesus-Hernandez, M., et al. Expanded GGGGCC hexanucleotide repeat in noncoding region of C9ORF72 causes chromosome 9p-linked FTD and ALS. Neuron 72, 245–256 (2011).

6. Renton, A.E., et al. A hexanucleotide repeat expansion in C9ORF72 is the cause of chromosome 9p21-linked ALS-FTD. Neuron 72, 257–268 (2011).

7. Ilieva, H., Polymenidou, M. & Cleveland, D.W. Non-cell autonomous toxicity in neurodegenerative disorders: ALS and beyond. J Cell Biol 187, 761–772 (2009).

8. Yamanaka, K. & Komine, O. The multi-dimensional roles of astrocytes in ALS. Neurosci Res 126, 31–38 (2018).

9. Schmidt, E.R., Pasterkamp, R.J. & van den Berg, L.H. Axon guidance proteins: novel therapeutic targets for ALS? Prog Neurobiol 88, 286–301 (2009).

10. Klein, R. Bidirectional modulation of synaptic functions by Eph/ephrin signaling. Nat Neurosci 12, 15–20 (2009).

11. Pasquale, E.B. Eph-ephrin bidirectional signaling in physiology and disease. Cell 133, 38–52 (2008).

12. Murai, K.K., Nguyen, L.N., Irie, F., Yamaguchi, Y. & Pasquale, E.B. Control of hippocampal dendritic spine morphology through ephrin-A3/EphA4 signaling. Nat Neurosci 6, 153–160 (2003).

13. Ohta, K., et al. The receptor tyrosine kinase, Cek8, is transiently expressed on subtypes of motoneurons in the spinal cord during development. Mech Dev 54, 59–69 (1996).

14. Kandouz, M. Dying to communicate: apoptotic functions of Eph/Ephrin proteins. Apoptosis 23, 265–289 (2018).

15. Li, J., et al. Inhibition of EphA4 signaling after ischemia-reperfusion reduces apoptosis of CA1 pyramidal neurons. Neurosci Lett 518, 92–95 (2012).

16. Van Hoecke, A., et al. EPHA4 is a disease modifier of amyotrophic lateral sclerosis in animal models and in humans. Nat Med 18, 1418–1422 (2012).

17. Rue, L., et al. Reduction of ephrin-A5 aggravates disease progression in amyotrophic lateral sclerosis. Acta Neuropathol Commun 7, 114 (2019).

18. Ling, K.K., et al. Antisense-mediated reduction of EphA4 in the adult CNS does not improve the function of mice with amyotrophic lateral sclerosis. Neurobiol Dis 114, 174–183 (2018).

19. Rue, L., et al. Reducing EphA4 before disease onset does not affect survival in a mouse model of Amyotrophic Lateral Sclerosis. Sci Rep 9, 14112 (2019).

20. Zhao, J., Cooper, L.T., Boyd, A.W. & Bartlett, P.F. Decreased signalling of EphA4 improves functional performance and motor neuron survival in the SOD1(G93A) ALS mouse model. Sci Rep 8, 11393 (2018).

21. Bundesen, L.Q., Scheel, T.A., Bregman, B.S. & Kromer, L.F. Ephrin-B2 and EphB2 regulation of astrocyte-meningeal fibroblast interactions in response to spinal cord lesions in adult rats. J Neurosci 23, 7789–7800 (2003).

22. Fabes, J., et al. Accumulation of the inhibitory receptor EphA4 may prevent regeneration of corticospinal tract axons following lesion. Eur J Neurosci 23, 1721–1730 (2006).

23. Mitsumoto, H., Chad, D.A. & Pioro, E.P. Amyotrophic lateral sclerosis, (F.A. Davis, Philadelphia, 1998).

24. McClelland, A.C., Sheffler-Collins, S.I., Kayser, M.S. & Dalva, M.B. Ephrin-B1 and ephrin-B2 mediate EphB-dependent presynaptic development via syntenin-1. Proc Natl Acad Sci U S A 106, 20487–20492 (2009).

25. Lepore, A.C., et al. Focal transplantation-based astrocyte replacement is neuroprotective in a model of motor neuron disease. Nat Neurosci 11, 1294–1301 (2008).

26. Nicaise, C., et al. Early phrenic motor neuron loss and transient respiratory abnormalities following unilateral cervical spinal cord contusion. J Neurotrauma (2013).

27. Nicaise, C., et al. Phrenic motor neuron degeneration compromises phrenic axonal circuitry and diaphragm activity in a unilateral cervical contusion model of spinal cord injury. Exp Neurol 235, 539–552 (2012).

28. Nicaise, C., et al. Degeneration of phrenic motor neurons induces long-term diaphragm deficits following mid-cervical spinal contusion in mice. J Neurotrauma 29, 2748–2760 (2012).

29. Lepore, A.C., et al. Human glial-restricted progenitor transplantation into cervical spinal cord of the SOD1^G93A^ mouse model of ALS. PLoS One 6(2011).

30. Lepore, A.C., et al. Peripheral hyperstimulation alters site of disease onset and course in SOD1 rats. Neurobiol Dis 39, 252–264 (2010).

31. Lepore, A.C., et al. Intraparenchymal spinal cord delivery of adeno-associated virus IGF-1 is protective in the SOD1G93A model of ALS. Brain Res 1185, 256–265 (2007).

32. Tsuda, H.H., S.M.; Yang, Y.; Tong, C.; Lin, Y.Q.; Mohan, K.; Haueter, C.; Zoghbi, A.; Harati, Y.; Kwan, J.; Miller, M.A.; Bellen, H.J. The amyotrophic lateral sclerosis 8 protein VAPB is cleaved, secreted, and acts as a ligand for Eph receptors. Cell 133(6), 963–977 (2008).

33. Warren, P.M. & Alilain, W.J. The challenges of respiratory motor system recovery following cervical spinal cord injury. Prog Brain Res 212, 173–220 (2014).

34. Lepore, A.C. & Fischer, I. Lineage-restricted neural precursors survive, migrate, and differentiate following transplantation into the injured adult spinal cord. Exp Neurol 194, 230–242 (2005).

35. Llado, J., et al. Degeneration of respiratory motor neurons in the SOD1 G93A transgenic rat model of ALS. Neurobiol Dis 21, 110–118 (2006).

36. Yamanaka, K., et al. Astrocytes as determinants of disease progression in inherited amyotrophic lateral sclerosis. Nat Neurosci 11, 251–253 (2008).

37. Wright, M.C., Cho, W.J. & Son, Y.J. Distinct patterns of motor nerve terminal sprouting induced by ciliary neurotrophic factor vs. botulinum toxin. J Comp Neurol 504, 1–16 (2007).

38. Wright, M.C., et al. Distinct muscarinic acetylcholine receptor subtypes contribute to stability and growth, but not compensatory plasticity, of neuromuscular synapses. J Neurosci 29, 14942–14955 (2009).

39. Wright, M.C. & Son, Y.J. Ciliary neurotrophic factor is not required for terminal sprouting and compensatory reinnervation of neuromuscular synapses: re-evaluation of CNTF null mice. Experimental neurology 205, 437–448 (2007).

40. Li, K., et al. GLT1 overexpression in SOD1(G93A) mouse cervical spinal cord does not preserve diaphragm function or extend disease. Neurobiol Dis 78, 12–23 (2015).

41. Martin, M., Li, K., Wright, M.C. & Lepore, A.C. Functional and morphological assessment of diaphragm innervation by phrenic motor neurons. J Vis Exp, e52605 (2015).

42. Tyzack, G.E.H., C.E.; Sibley, C.R.; Cymes, T.; Forostyak, S.; Carlino, G.; Meyer, I.F.; Schiavo, G.; Zhang, S.C.; Gibbons, G.M.; Newcombe, J.; Patani, R.; Lakatos, A. A neuroprotective astrocyte state is induced by neuronal signal EphB1 but fails in ALS models. Nat Commun 8(1), 1164 (2017).

43. Zhao, J.B., A.W.; Bartlett, P.F. The identification of a novel isoform of EphA4 and ITS expression in SOD1G93A mice. Neuroscience 347, 11–21 (2017).

44. Rothstein, J.D., Van Kammen, M., Levey, A.I., Martin, L.J. & Kuncl, R.W. Selective loss of glial glutamate transporter GLT-1 in amyotrophic lateral sclerosis. Ann Neurol 38, 73–84 (1995).

45. Phatnani, H.P.G., P.; Friedman, B.A.; Carrasco, M.A.; Muratet, M.; O’Keeffe, S.; Nwakeze, C.; Pauli-Behn, F.; Newberry, K.M.; Meadows, S.K.; Tapia, J.C.; Myers, R.M.; Maniatis, T. Intricate interplay between astrocytes and motor neurons in ALS. Proc Natl Acad Sci U S A 110(8), E756–765 (2013).

46. Cassina, P.C., A.; Pehar, M.; Castellanos, R.; Gandelman, M.; de León, A.; Robinson, K.M.; Mason, R.P.; Beckman, J.S.; Barbeito, L.; Radi, R. Mitochondrial dysfunction in SOD1G93A- bearing astrocytes promotes motor neuron degeneration: prevention by mitochondrial-targeted antioxidants. J Neurosci 28(16), 4115–4122 (2008).

47. Flanagan, J.G. & Vanderhaeghen, P. The ephrins and Eph receptors in neural development. Annu Rev Neurosci 21, 309–345 (1998).

48. Wu, B., et al. Potent and Selective EphA4 Agonists for the Treatment of ALS. Cell Chem Biol 24, 293–305 (2017).

49. Qin, H., Lim, L.Z. & Song, J. Dynamic principle for designing antagonistic/agonistic molecules for EphA4 receptor, the only known ALS modifier. ACS Chem Biol 10, 372–378 (2015).

50. Schoonaert, L., et al. Identification and characterization of Nanobodies targeting the EphA4 receptor. J Biol Chem 292, 11452–11465 (2017).

51. Henderson, N.T. & Dalva, M.B. EphBs and ephrin-Bs: Trans-synaptic organizers of synapse development and function. Mol Cell Neurosci 91, 108–121 (2018).

52. Hanamura, K., et al. Extracellular phosphorylation of a receptor tyrosine kinase controls synaptic localization of NMDA receptors and regulates pathological pain. PLoS Biol 15, e2002457 (2017).

53. Nolt, M.J., et al. EphB controls NMDA receptor function and synaptic targeting in a subunit-specific manner. J Neurosci 31, 5353–5364 (2011).

54. Alkaslasi, M.R., et al. Single nucleus RNA-sequencing defines unexpected diversity of cholinergic neuron types in the adult mouse spinal cord. Nat Commun 12, 2471 (2021).

55. Blum, J.A., et al. Single-cell transcriptomic analysis of the adult mouse spinal cord reveals molecular diversity of autonomic and skeletal motor neurons. Nat Neurosci 24, 572–583 (2021).

56. Van Harten, A.C.M., Phatnani, H. & Przedborski, S. Non-cell-autonomous pathogenic mechanisms in amyotrophic lateral sclerosis. Trends Neurosci 44, 658–668 (2021).

57. Mirian, A., et al. Breached Barriers: A Scoping Review of Blood-Central Nervous System Barrier Pathology in Amyotrophic Lateral Sclerosis. Front Cell Neurosci 16, 851563 (2022).

58. Makinen, P.I., et al. Stable RNA interference: comparison of U6 and H1 promoters in endothelial cells and in mouse brain. J Gene Med 8, 433–441 (2006).

59. Falnikar, A., Hala, T.J., Poulsen, D.J. & Lepore, A.C. GLT1 overexpression reverses established neuropathic pain-related behavior and attenuates chronic dorsal horn neuron activation following cervical spinal cord injury. Glia 64, 396–406 (2016).

60. Li, K., et al. Transplantation of glial progenitors that overexpress glutamate transporter GLT1 preserves diaphragm function following cervical SCI. Mol Ther 23, 533–548 (2015).

61. Li, K., et al. Overexpression of the astrocyte glutamate transporter GLT1 exacerbates phrenic motor neuron degeneration, diaphragm compromise, and forelimb motor dysfunction following cervical contusion spinal cord injury. J Neurosci 34, 7622–7638 (2014).

62. Lepore, A.C., et al. Selective ablation of proliferating astrocytes does not affect disease outcome in either acute or chronic models of motor neuron degeneration. Exp Neurol 211, 423–432 (2008).

63. Andersen, P.M., et al. Phenotypic heterogeneity in motor neuron disease patients with CuZn-superoxide dismutase mutations in Scandinavia. Brain 120 (Pt 10), 1723–1737 (1997).

64. Radunovic, A. & Leigh, P.N. Cu/Zn superoxide dismutase gene mutations in amyotrophic lateral sclerosis: correlation between genotype and clinical features. J Neurol Neurosurg Psychiatry 61, 565–572 (1996).

65. Goulao, M., et al. Astrocyte progenitor transplantation promotes regeneration of bulbospinal respiratory axons, recovery of diaphragm function, and a reduced macrophage response following cervical spinal cord injury. Glia 67, 452–466 (2019).

66. Lepore, A.C. Intraspinal cell transplantation for targeting cervical ventral horn in amyotrophic lateral sclerosis and traumatic spinal cord injury. J Vis Exp (2011).

67. Li, K., et al. Human iPS cell-derived astrocyte transplants preserve respiratory function after spinal cord injury. Exp Neurol 271, 479–492 (2015).

68. Cheng, L., et al. LAR inhibitory peptide promotes recovery of diaphragm function and multiple forms of respiratory neural circuit plasticity after cervical spinal cord injury. Neurobiology of Disease (In press).

69. Ghosh, B., et al. Local BDNF delivery to the injured cervical spinal cord using an engineered hydrogel enhances diaphragmatic respiratory function. J Neurosci 38, 5982–5995 (2018).

70. Ghosh, B., et al. A hydrogel engineered to deliver minocycline locally to the injured cervical spinal cord protects respiratory neural circuitry and preserves diaphragm function. Neurobiol Dis 127, 591–604 (2019).

